# Doxycycline interferes with tau amyloid aggregation abolishing its associated neuronal toxicity

**DOI:** 10.1101/2020.11.18.388561

**Authors:** Luciana Medina, Florencia González-Lizárraga, Antonio Dominguez-Meijide, Diego Ploper, Valeria Parrales, Sabrina Sequeira, Maria Sol Cima-Omori, Markus Zweckstetter, Elaine Del Bel, Patrick P. Michel, Tiago Fleming Outeiro, Rita Raisman-Vozari, Rosana Chehín, Sergio B. Socias

## Abstract

Tauopathies are neurodegenerative disorders with increasing incidence and still without cure. The extensive time required for development and approval of novel therapeutics highlights the need for testing and repurposing known safe molecules. Since doxycycline impacts α-synuclein aggregation and toxicity, herein we tested its effect on tau. We found that doxycycline reduces amyloid aggregation of the different isoforms of tau protein in a dose-dependent manner, remodeling the resultant species. Furthermore, doxycycline interacts with tau microtubule-binding domain preventing its aggregation. In a cell free system doxycycline also prevents tau seeding and in cell culture reduces toxicity of tau aggregates. Overall, our results expand the spectrum of action of doxycycline against aggregation-prone proteins, opening novel perspectives for its repurposing as a disease-modifying drug for tauopathies.

---

Tauopathies are a group of neurodegenerative diseases characterized by clinical heterogeneity and progressive deposition of amyloid aggregates of abnormally hyper-phosphorylated tau protein within specific brain regions [1]. In healthy neurons, tau is the major microtubule associated protein (MAP), and plays a crucial role in regulating its dynamics, concomitant axonal transport and neurite outgrowth [2].

From a biochemical point of view, tau is an intrinsically disordered protein (IDP) with an N-terminal “projection domain” that projects away from microtubules [3]; and a positively charged C-terminal domain that binds tubulin and promotes self-assembly [4–6]. Six tau isoforms are expressed in the human brain as a consequence of the alternative splicing. This in turn, leads to the expression of the tau proteins 0N3R, 1N3R, 2N3R, 0N4R, 1N4R and 2N4R, being N the 29 amino acid N-terminal inserts and R the microtubule binding repeats [2].

In a physiological context, tau is regulated by site-specific phosphorylation. However, in pathological conditions, abnormal phosphorylation and aggregation lead to the formation of tau amyloid aggregates called paired helical filaments (PHFs), which ultimately lead to the build-up of cytoplasmic neurofibrillary tangles (NFT) [7]. Although the topographical distribution patterns of the lesions that contain NFTs correlate with the clinical progression of tauopathies such as Alzheimer’s disease (AD) [8], the mechanisms of tau-associated neurodegeneration remain unclear. Likewise, despite the fact that the relationship between tau phosphorylation and aggregation was found to play a central role in the transition from its native state to the pathological form [9–13], the relative contribution of each process to disease etiology and progression is poorly understood.

Unfortunately, albeit tremendous efforts and massive investments, therapies capable of preventing, halting, or at least slowing the progression of tau-associated disorders are not available [14]. In this context, the aging of the human population, the main risk factor for AD and other neurodegenerative diseases, threatens to burden healthcare systems worldwide [15]. Therefore, it is urgent to develop disease-modifying therapies, a process that is proving extremely difficult and takes many years from bench to bedside.

An alternative strategy that would significantly reduce time and costs is “drug repurposing”, which involves the use of pre-existing and approved drugs for new indications [16]. Compelling evidence show that many antibiotics, *i.e*. minocycline and doxycycline [17], aside from their antimicrobial action, are capable of halting the noxious amyloid aggregation process of disease-associated proteins [18–22]. Therefore, this positions them as promising alternatives for the development of efficient therapies against neurodegenerative disorders. In this regard, we recently demonstrated the ability of doxycycline to reshape early oligomers of α-synuclein, preventing the buildup of pathogenic species and ultimately redirecting the process towards non-toxic off-pathway oligomers [23].

In the present work, we studied the effect of doxycycline on tau amyloid aggregation pathway and its associated toxicity. By using heparin-induced 2N4R tau fibrillization as well as the 4R truncated species self-aggregation, we assessed the capacity of doxycycline to hinder the tau amyloid pathway. Our results suggest the relevance of tau microtubule binding domain in doxycycline:tau interaction. Additionally, doxycycline also halted the ability of tau seeds to recruit monomers, which is essential for the progression of the pathology. By using cell culture, we also demonstrate the ability of this tetracycline to abate the toxicity related with tau-aggregated species. Taking together these results, with the well-known brain bioavailability, safety, anti-inflammatory and antioxidant abilities of doxycycline, endorse this old drug as an ideal compound to be repurposed for tauopathies.

## Materials and methods

### Preparation of heterologous 2N4R tau

Expression and purification of recombinant human tau was performed as previously described by Barghorn et al. [24] using the plasmid tau/pET29b (Addgene, #16316) and *E. coli* BL21 [DE3]. Briefly, the purity of the protein was assessed by SDS-PAGE. Monomeric tau stock solutions were prepared in 40 mM Tris-HCl, 20 mM MgCl_2_ pH 7.5 with 0.05% DTT (freshly added). Prior to measurements, protein solutions were centrifuged for 30 min at 12000 x *g* and filtered with a 0.22 μm filter (Millex-GV. Millipore). Protein concentration was determined by the measurement of absorbance at 280 nm using extinction coefficient ɛ = 7700 cm^−1^.M^−1^. Tau/pET29b was a gift from Peter Klein (Addgene plasmid # 16316 ; http://n2t.net/addgene:16316 ; RRID:Addgene_16316) [25].

### Protein aggregation

Monomeric tau (22 μM) in 40 mM Tris-HCl, 20 mM MgCl_2_ pH 7.5 and fresh 0.05% DTT, was incubated in a Thermomixer Comfort® (Eppendorf) at 37°C under orbital agitation at 600 rpm; using 0.2 mg/ml heparin, in the absence or presence of doxycycline.

### Thioflavin T assay

Formation of cross-β structure during tau aggregation was followed by addition of Thioflavin T (ThT) fluorescent probe on aliquots withdrawn from the incubation mixture at different times, according to LeVine[26,27]. Changes in the emission fluorescence spectra with the excitation wavelength set at 450 nm were monitored using an ISS (Champaign, IL) PC1 spectrofluorometer. Doxycycline dose-response assay on tau aggregation was fitted to the equation:

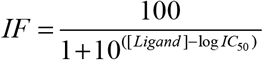

where IF is the normalized fluorescence intensity, [Ligand] is doxycycline concentration, and IC_50_ is the concentration at which aggregation is inhibited at a 50%.

### Bis-ANS fluorescent assay

Tau was aggregated as described above in the absence or presence of doxycycline. Aliquots were taken at 0 and 24 h incubation, and bis (1-Anilinonaphthalene-8-Sulfonic Acid) also known as bis-ANS was added to a final concentration of 5 μM. Bis-ANS was excited at 395 nm and fluorescence emission was measured from 410 nm to 610 nm.

### Infrared Spectroscopy

Samples of tau (100 μM), tau:heparin (100 μM:0.8 mg/ml) and tau:heparin:doxycycline (100 μM:0.8 mg/ml:400 μM) in buffer-D_2_O 40 mM Tris-HCl, 20 mM MgCl_2_, pH 7.5, 0.05% DTT, were collected after 24 h incubation at 37°C under orbital agitation (600 rpm) and assembled between two CaF_2_ windows with a path length of 50 nm in a thermostated cell. The spectra were recorded in a Nicolet 5700 spectrometer (Thermo Nicolet, Madison, WI) equipped with a DTGS detector as described by Arrondo et al. [28]. The sample chamber was permanently purged with dry air. The buffer spectra were subtracted from that of the solution at the same temperature in order to eliminate the D_2_O contribution in the Amide I’ region to get a flat baseline between 1,900 and 1,700 cm^−1^. After solvent subtraction, quantitative information on protein structure was obtained through deconvolution and derivation of the Amide I’ band into its constituents [28]. To obtain the relative contribution of each component, an iterative process based on minimal square was performed by using curve fit under SpectraCalc software. The mathematical solution of the decomposition may not be unique, but if restrictions are imposed such as the maintenance of the initial band positions in an interval of 1 cm^−1^, the preservation of the bandwidth within the expected limits, or the agreement with theoretical boundaries or predictions, the result becomes, in practice, unique.

### Transmission Electron Microscopy

4 μl of 22 μM tau aggregation samples were adsorbed onto glow-discharged 200 mesh formvar carbon coated copper grids (Electron Microscopy Sciences) and stained with aqueous uranyLess (Electron Microscopy Sciences). After washing, excess liquid was removed, and grids were allowed to air dry. Transmission electron microscopy micrographs were collected using a Hitachi 7700 transmission electron microscope.

### Protease resistance assay

Tau, tau:heparin and tau.heparin:doxycycline samples obtained after 168 h of incubation were used as substrate of proteases in the digestion assay. Reactions were carried out mixing tau samples with proteinase K (152g/ml) or trypsin (0.0125 %), for 30 minutes at 37°C in buffer 40 mM Tris-HCl, 20 mM MgCl_2_ pH 7.5 with 0.05% DTT. After incubation, the enzymes were inactivated with PMSF 1mM and subsequently loading buffer was added to each sample for 12% Tris-Glycine SDS-PAGE gel. The gel was stained with colloidal Coomasie Blue incubating with gentle stirring throughout the night. Photographs were taken and the gels were analyzed using Image J 1.47v software (National Institutes of Health, EUA), to obtain the densitometric profiles of the bands of each street [29].

### Real time quaking induced conversion (RT-QuiC)

RT-QuiC reactions were performed in black 96-well plates (COSTAR, Corning Incorporated) in which 100 μl of the reaction mixtures were loaded. Mix were prepared to the following final concentrations: 150 mM NaCl, 1 mM EDTA, 10 μM Thioflavin T, 70 μM SDS, and 0.5 μM of the monomeric microtubule binding domain of 2N4R tau (K18 peptide) in PBS buffer (pH= 7.1). Doxycycline was added at the indicated final concentrations. Plates were covered with sealing tape and incubated at 41°C in a plate reader (Infinite M200 fluorescence plate reader, TECAN) with intermittent cycles of one-minute orbital shaking at 432 rpm followed by 2 min incubation and 1 min pause to measure the fluorescence intensity at 480 nm. Three replicates of each sample were measured for 250 amplification cycles.

### Seeding assay

Samples of tau:heparin (100 μM:0.8 mg/ml) in buffer 40 mM Tris-HCl, 20 mM MgCl_2_, pH 7.5, 0.05% DTT, were incubated for 24 h in a Thermomixer Comfort® (Eppendorf) at 37°C under orbital agitation at 600 rpm. These species were diluted 1/10 when harvested and further incubated with 22 μM of fresh monomers in the absence or presence of 100 μM doxycycline. Seeding aggregation was followed using ThT fluorescent probe. Aliquots were taken from the seeding aggregation after 168 h and mixed with ThT, according to LeVine [26,27], as previously described, using FluoroMax-4 Spectrofluorometer.

### Human neuroblastoma cell culture and cytotoxicity assay

SH-SY5Y cells were grown in DMEM supplemented with 10% fetal bovine serum (FBS) and 1% penicillin/streptomycin (PS), at 37°C and 5%CO_2_. For cell viability assay cells were seeded in 96 wells plates at 15000 cells/well and maintained in 100 μl of DMEM supplemented with 2% FBS and 1% PS for 24 h at 37°C. Afterwards, cells were treated with a 25 μl aliquot of pre-incubated (37°C, 6000 rpm 16 h) tau, tau:heparin, tau:heparin:doxycyline and incubated for 24 h at 37°C 5% CO_2_. To determine cell viability, the colorimetric MTT metabolic activity assay was used as previously described by Mosmann [30]. All experiments were performed in sextuplicate, and the relative cell viability (%) was expressed as a percentage relative to the untreated cell control.

### Statistical analyses

All data were obtained from at least three independent experiments and expressed as mean ± SD. Multiple-group comparisons were performed with one-way ANOVA and t-test. Differences were considered as statistically significant at p < 0.05. Statistical analyses were carried out with GraphPad Prism 5 (San Diego, California, USA).

## RESULTS

### Doxycycline hinders tau amyloid fibril formation yielding novel species

The ability of doxycycline to interfere with 2N4R tau aggregation was studied in the presence of heparin, a classical model system which induces the formation of tau fibrillary elements similar to NFT [10,13]. In agreement with previous reports, we found that heparin efficiently triggered full-length tau amyloid aggregation as monitored by Thioflavin T (ThT) fluorescence emission. Doxycycline inhibited tau aggregation in a dose-dependent manner (Fig. 1A), showing an IC_50_ of 29 μM (Fig. 1B). Since doxycycline exerted optimal inhibition of tau amyloid aggregation at 100 μM, all experiments henceforth were performed at this concentration. Kinetics of heparin–induced tau aggregation showed a classical sigmoidal behavior (Fig. 1C), characteristic of the nucleation-polymerization process as previously described [31–33]. The short lag phase was followed by an exponential increase that finally reached a plateau after 24 h of incubation. In the presence of doxycycline the system followed a hyperbolic curve, suggesting that early species formed in the presence of the tetracycline poisoned the cooperative characteristic of this aggregation process. Moreover, the amount of cross-β-containing species finally formed was significantly reduced in the presence of doxycycline, with a fluorescence steady state remaining stable up to 168 h (Fig. 1C). The different kinetics observed in the presence or absence of doxycycline suggests that distinct tau species were formed. To evaluate this hypothesis, we compared the hydrophobic patches exposed to solvent of these species. For this, we studied the interaction of bis-ANS with species prepared with or without doxycycline (Fig. 1D). Basal bis-ANS emission spectrum of tau increased when heparin was added to trigger amyloid aggregation, reflecting a gain of hydrophobic surfaces exposure during the process. On the contrary, in the presence of doxycycline, the fluorescence intensity diminished significantly (61%) (Fig. 1D). These results indicate that these species were structurally different and the presence of doxycycline led to less hydrophobic residues exposed.

**Fig. 1.**
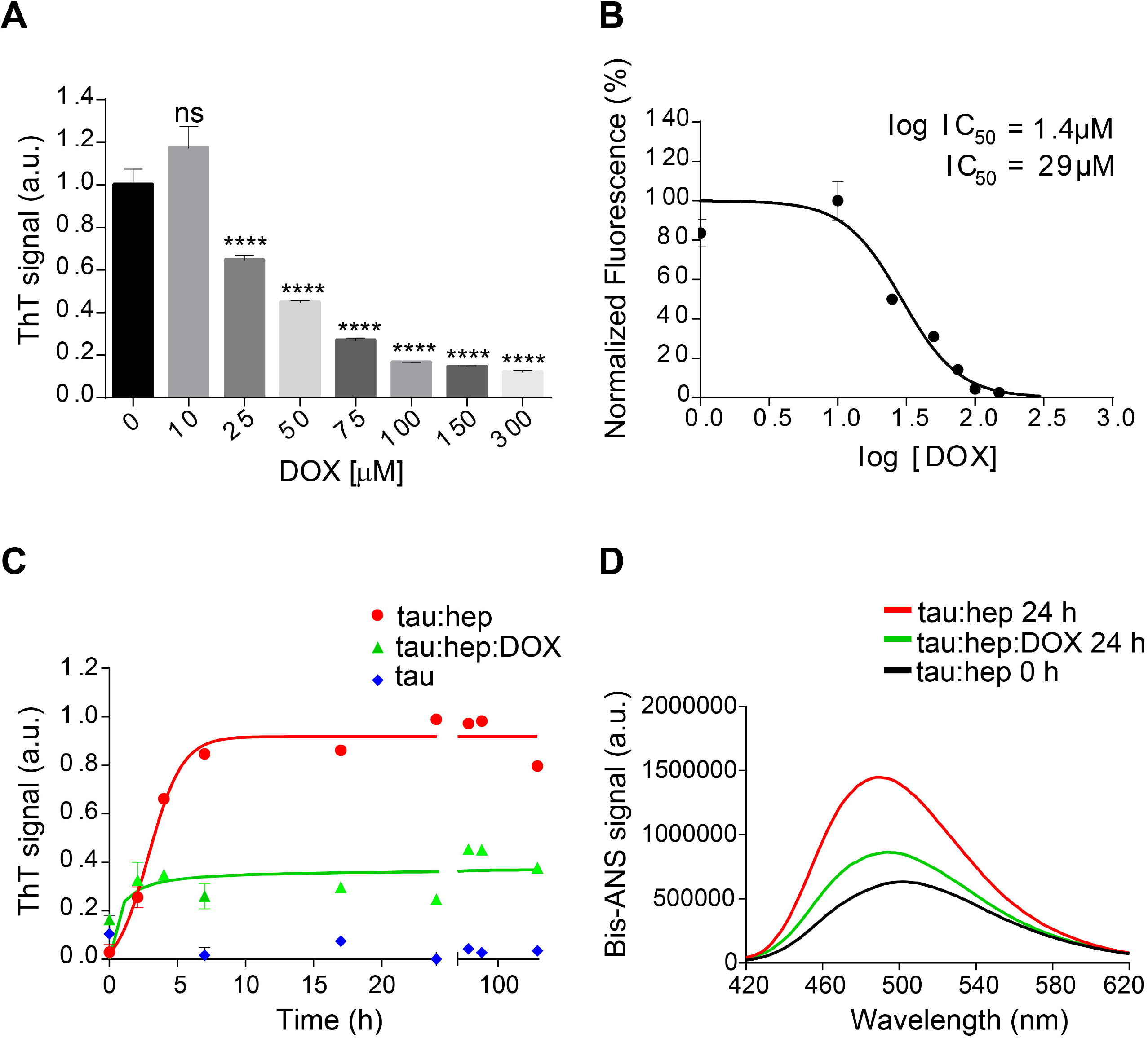
Doxycycline affects tau canonical amyloid aggregation. **A.** Fluorescence emission intensity of 25 μM of thioflavin T in a solution containing samples of 22 μM tau, 0.2 mg/ml heparin and 10 μM, 25 μM, 75 μM, 100 μM, 150 μM or 300 μM of doxycycline. Samples were incubated at 37°C under orbital agitation and aggregation was assayed after 24 h by ThT fluorescence emission. n = 3 ns: not significant. * *p* ≤ 0.05; ** *p* ≤ 0.01; *** *p* ≤ 0.001; **** *p* ≤ 0.0001. Error bars represent SD. **B.** Dose-dependent response of ThT fluorescence of tau:heparin solution (tau 22 μM; 0.2 mg/ml heparin) incubated at 37°C under orbital agitation for 24 h, in the presence of different concentrations of doxycycline. IC_50_ values are mean values of three independent determinations. The data fitting is described in Methods section. **C.** Fluorescence emission intensity of 25 μM thioflavin T in a solution containing samples of tau 22 μM; 0.2 mg/ml heparin; and 100 μM of doxycycline. Samples were incubated at 37°C under orbital agitation and aggregation was assayed by ThT fluorescence emission. **D.** Bis-ANS fluorescence signal of tau:heparin solution incubated 0 h (black line) and 24 h at 37°C under orbital agitation in the absence (red line) or presence of doxycycline (green line).

### Doxycycline affects tau seeding ability and toxicity

Considering that hydrophobicity is one of the primary driving forces behind protein self-assembly processes [34,35], and that brain-derived tau oligomeric species that can spread the pathology have affinity for bis-ANS [36], we assessed the ability of doxycycline at halting the pro-aggregating properties of heparin-induced tau aggregates on monomeric species. For this, tau seeds were produced by incubating monomeric species with heparin at 37°C under orbital agitation for 24 h. Then, aliquots of this solution were added to fresh tau samples (deprived of heparin) with and without 100 μM of doxycycline. Only in the absence of the tetracycline, the seeding effect of pre-aggregated tau on the monomeric protein could be observed (Fig. 2A). In the absence of seeds, monomeric tau did not evolve into amyloid species, as indicated by reduced ThT signal. As an internal control, heparin at the residual concentration present in seed aliquots, was added to monomeric tau and the mix incubated in the same experimental conditions. Upon incubation, no amyloid aggregation was observed according to ThT signal (Data not shown).

**Fig. 2.**
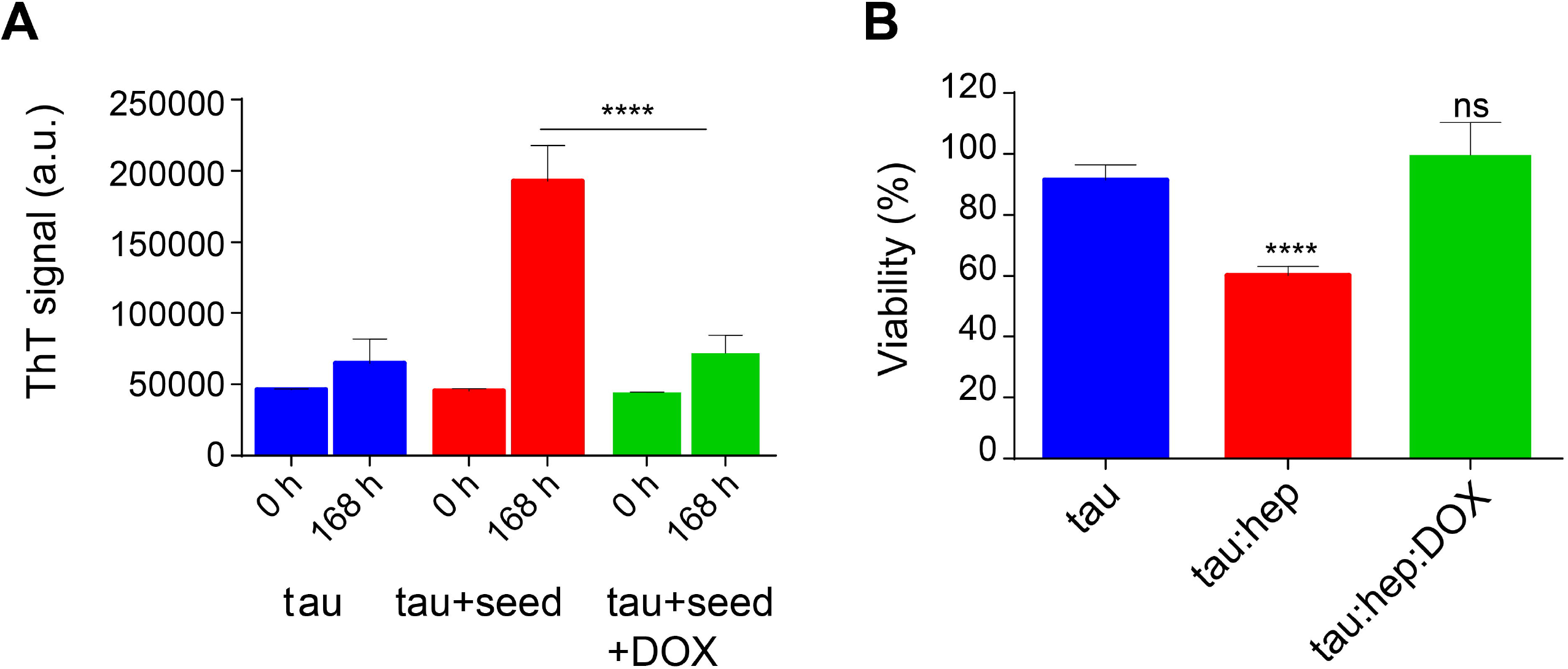
Doxycycline affects tau seeding ability and toxicity. **A.** Effect of doxycycline on the seeding capability of pre-aggregated tau species. Pre-incubated tau species were obtained by incubating a tau:heparin solution (tau 22 μM; 0.2 mg/ml heparin) 24 h at 37°C under orbital agitation. 25 μl aliquots of this solution were used as seeds and added to fresh monomeric tau (22 μM) in the absence or presence of doxycycline 100 μM and incubated 168 h at 37°C under orbital agitation. **B.** SH-SY5Y cells were treated with aggregated species of tau incubated in the presence of heparin, with or without doxycycline, and harvested after 16 h incubation. Viability was measured using an MTT assay and expressed as a percentage relative to the untreated cells. Statistical analyses for both experiments were performed using the ANOVA test. n = 3 ns: not significant. * *p* ≤ 0.05; ** *p* ≤ 0.01; *** *p* ≤ 0.001; **** *p* ≤ 0.0001. Error bars represent SD.

As our results showed that doxycycline remodeled tau aggregates affecting the formation of hydrophobic patches, we evaluated the ability of these species to bind and disrupt cellular membranes. For this, we analyzed cellular viability in a SH-SY5Y cell line using the MTT assay [30], which reflects the number of viable cells based on mitochondrial activity. SH-SY5Y cells were incubated with 25 μl aliquots of heparin-induced tau species prepared in the absence or presence of doxycycline and harvested after 16 h of incubation [37,38]. Cells were further incubated at 37°C for 24 h and their viability was assessed. Fig. 2B shows that, in good agreement with previous reports, heparin-induced tau oligomers harvested after 16 h of incubation in the absence of doxycycline led to a decrease of about 35 % in cell viability. On the contrary, the viability of cells treated with tau species prepared in the presence of doxycycline, showed no significant difference from control on MTT turnover, indicating that the tetracycline counteracts the gain of toxicity of the heparin-induced aggregation process (Fig. 2B). A solution of heparin was tested without significant differences from control (Data no shown).

In summary, Fig. 2 shows that the presence of doxycycline interfered with the seeding-ability of tau and rendered tau aggregates that were less toxic for cultured cells.

### Doxycycline diminished β-structuration of tau aggregates

The impact of doxycycline on the structure of heparin-induced tau aggregates was analyzed by using Fourier-Transform infrared spectroscopy (FTIR). Comparative analysis of the conformationally sensitive band Amide I’ (1,700– 1,600 cm^−1^ of the infrared spectrum) of aggregates were obtained in the presence and absence of doxycycline. The tau Amide I’ contour is centered at about 1,640 cm^−1^, which is typical for unfolded proteins [39,40]. Structural analysis performed by a curve fitting process (see Materials and Methods) allowed for band assignments according to previous reports [41], and showed close agreement with elements described by NMR [41]. Unstructured regions represented more than 40% of Amide I’. β–sheets, loops and turns were also detected (Fig. 3A and Table I) and their relative contributions were in accordance with previous reports [41]. The band centered at 1,619 cm^−1^, which has ambiguous assignments [42,43], was attributed to type II polyproline helixes (PPII) since tau has a proline-rich region, with a PPII conformation [44,45].

**Fig. 3.**
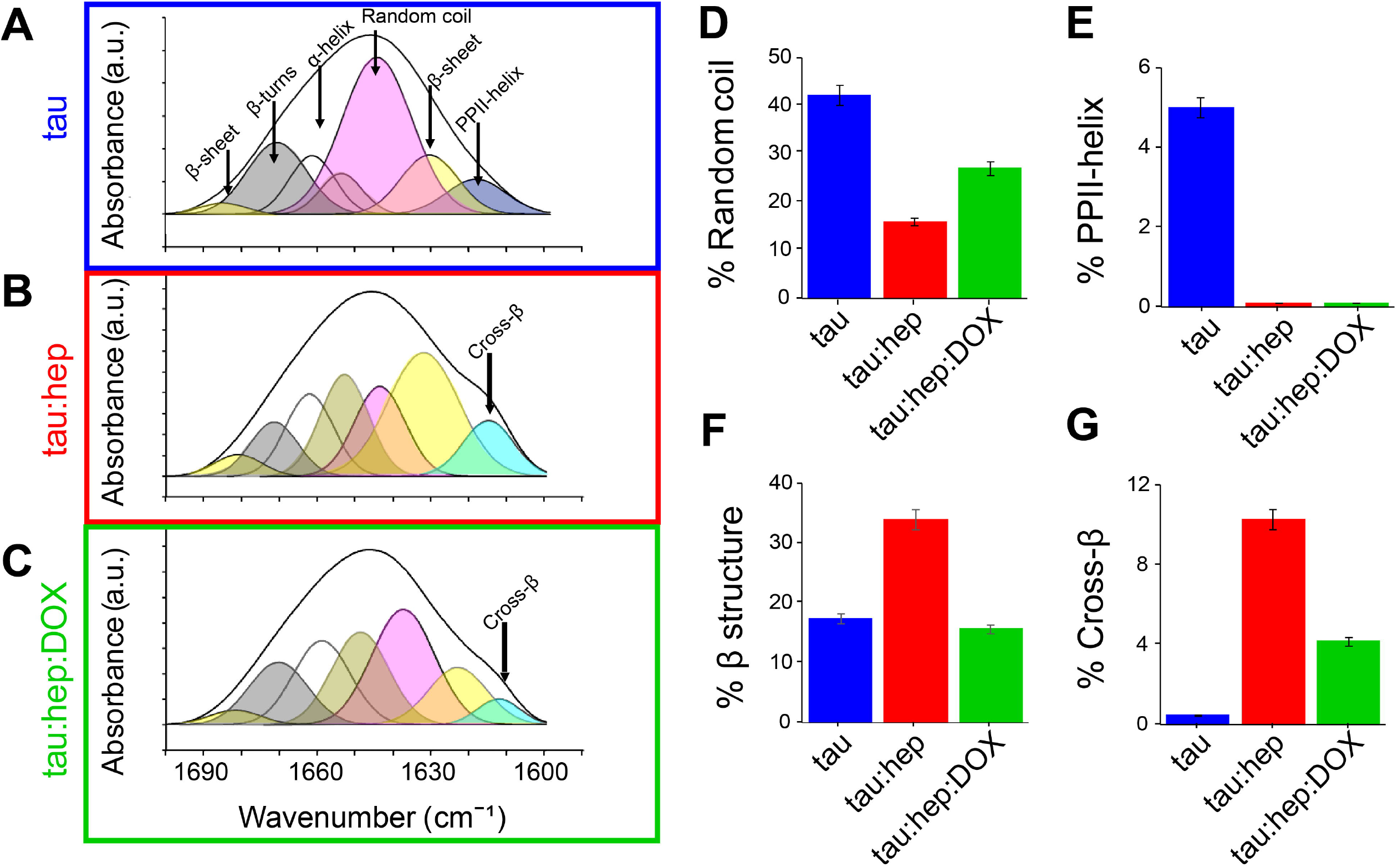
Doxycycline induces a different structural arrangement in tau aggregates. Tau FTIR Amide I’ curve fitting of **A)** 100 μM tau **B)**1 00 μM tau in the presence of heparin (0.8 mg/ml). **C)** 100 μM tau in the presence of heparin (0.8 mg/ml) and 100 μM doxycycline. Relative contribution of each component: **D)** Random coil, **E)** PPHII-helix, **F)** β structure, **G)** Cross-β. Error bars represent SD.

**Table 1.** FTIR-based evaluation of secondary structure content in different species of human tau incubated in the absence or presence of heparin and doxycycline for 24 h.

| Samples | Wavenumber (cm<sup>-1</sup>) | Relative Amide I' contribution | Assignments |
| --- | --- | --- | --- |
| tau | 1,610 | 1.2 | Side chains |
|  | 1,619 | 4.0 | PPII helix |
| | 1,631 | 15.9 | Low frequency β-sheet |
|  | 1,645 | 42.2 | Random coil |
| | 1,658 | 22.1 | α-helix + open loop |
| | 1,671 | 12.9 | β-turn |
| | 1,683 | 1.6 | High frequency β-sheet |
| tau:hep | 1,615 | 10.3 | Cross-β |
| | 1,632 | 30.6 | Low frequency β-sheet |
|  | 1,640 | 15.9 | Random coil |
| | 1,652 | 17.2 | α-helix |
|  | 1,662 | 13.6 | Open loop |
| | 1,672 | 8.9 | β-turn |
| | 1,681 | 3.3 | High frequency β-sheet |
| tau:hep:DOX | 1,614 | 4.2 | Cross-β |
| | 1,626 | 12.9 | Low frequency β-sheet |
|  | 1,637 | 27.0 | Random coil |
| | 1,649 | 20.3 | α-helix |
|  | 1,659 | 18.4 | Open loop |
| | 1,670 | 14.2 | β-turn |
| | 1,682 | 2.8 | High frequency β-sheet |

Tau Amide I’ spectrum remains stable after 24 h of incubation, which demonstrates the structural stability of the protein under our experimental conditions. However, in the presence of heparin, significant changes were detected (Fig. 3B). As expected according with ThT spectroscopy studies (Fig. 1C), a band located at around 1,614 cm^−1^ assignable to cross-β structure emerges, together with a significant increase in β-sheet structure. This β-structuration occurred at the expense of the random coil (Fig. 3B, D and Table I). The PPII-helix contribution was undetectable (Fig. 3F) even using high derivative or deconvolution factors.

The presence of doxycycline significantly diminished the 1,614 cm^−1^ band contribution, suggesting that this tetracycline hindered the heparin-induced gain of cross-β structures associated with amyloid aggregation of tau (Fig. 3C and G). Moreover, the overall β-structuration of the protein was decreased (curves shaded in yellow and light blue) (Fig. 3F), while non-structured regions (curve shaded in pink) remained more conserved (Fig. 3D).

### Doxycycline interferes with heparin-induced tau PHF-like formation

To evaluate whether doxycycline could affect heparin-induced full length (2N4R) tau aggregation, we used transmission electron microscopy (TEM). Samples of monomeric and aggregated tau species incubated with or without doxycycline and harvested after 168 h were analyzed (Fig. 4A). ThT fluorescence and infrared spectroscopy data showed that monomeric tau had no detectable assembly after 168 h incubation. In the presence of heparin, tau aggregated into long fibrils evoking PHF elements, as previously reported [10] (Fig. 4B). On the contrary, in the presence of doxycycline, the obtained aggregates lacked these characteristic elements. Instead, a mixture of oligomers and short fibrils abounded in all observed fields (Fig. 4C).

**Fig. 4.**
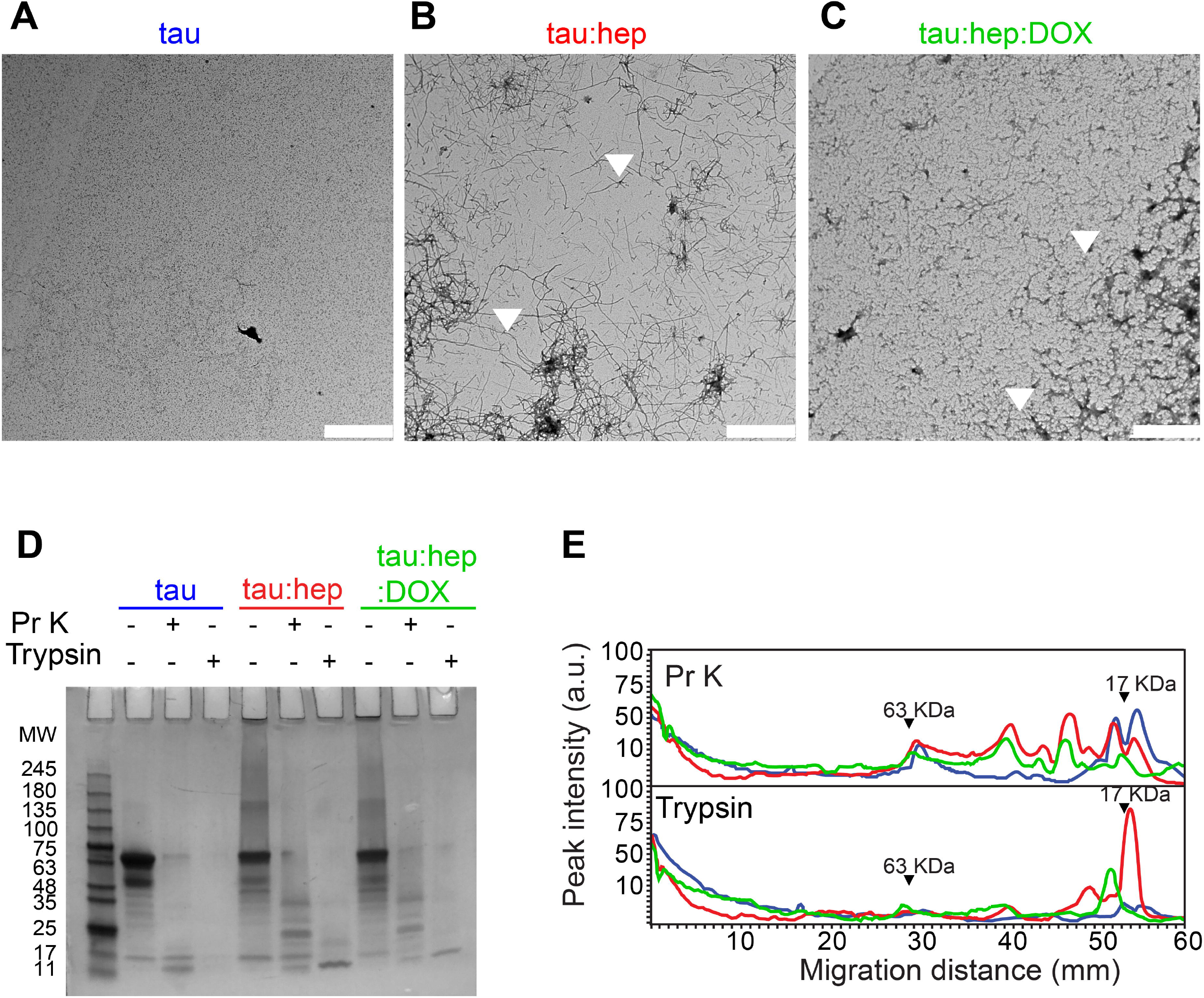
Doxycycline induces different quaternary structure arrangement of PHF-like fibrils. **A, B and C.** Transmission electron microscopy (TEM) of different 2N4R tau samples incubated at 37°C under orbital agitation and harvested after 168 h of incubation. Scale bar corresponds to 2 μm. **D.** Partial digestion profile of tau samples incubated in the same conditions as A, treated and not treated with 1 μg/ml proteinase K and with 0.0125% trypsin. Digestion products were resolved in a 12% tris-glycine gel stained with colloidal Coomasie Blue. Molecular weight marker in kDa. **E.** Densitometric analysis of the SDS-PAGE gel B were performed by using Image J 1.47v software.

Protease resistance profiles of full-length tau prepared in the absence or presence of doxycycline is depicted in Fig. 4D and E. Samples of monomeric tau as well as heparin-induced fibrils prepared in the absence or presence of doxycycline were harvested after 168 h of incubation and treated with proteinase K or trypsin. The trypsin limited digestion of tau fibrillar species results in a resistant core that comprises approximately the second half of R1, R2, R3 and the first half of R4 [46]. The removal of the fuzzy coat from the PHF left the fibril core exposed which was retained in the stacking gel as previously reported [47]. Digestion products were resolved on a 12% Tris-Glycine SDS-PAGE gel (Fig. 4D), and the corresponding densitometric analysis are shown in Fig. 4E. The data revealed that monomeric tau was digested into peptides of less than 11 kDa, while heparin-induced tau fibrils exhibited fragments resistant to both proteases, excised from the fibril core. In presence of doxycycline the SDS-PAGE and the densitogram showed a significant change in the digestion profile for both proteases, whose digestion products were enriched in low molecular weight elements. This data suggests that the doxycycline interacts with the microtubule-binding region, inducing novel conformational arrangements and exposing sites previously inaccessible to protease cleavage.

### Doxycycline interacts with tau microtubule binding domains inhibiting its self-aggregation

To confirm the hypothesis that doxycycline interacts with the microtubule-binding region of tau, we evaluated the effect of this tetracycline on the aggregation of a 4R truncated tau (K18 peptide), which can reach the fibrillary state without an inducer[48]. In order to evaluate the effect of different doxycycline concentrations on K18 peptide aggregation we used the real time quaking induced complementation (RT-QuiC) assay [49]. The results obtained showed that ThT fluorescence intensity values decreased at higher concentrations of doxycycline (Fig. 5A), abolishing the fluorescent signal at 70 μM. Furthermore, we observed a significantly lower K18 monomer incorporation rate per cycle in samples treated with 1, 10 and 70 μM doxycycline (Fig. 5B). Our results showed that doxycycline inhibited in a dose dependent manner K18 peptide amyloid aggregation, by blocking monomer incorporation to the growing fibrils. We suggest that doxycycline binds to the microtubule-binding region, considered the core of tau fibrils [50], disrupting the intermolecular interactions of tau at the level of these key repetitions in the process of amyloid aggregation.

**Fig. 5.**
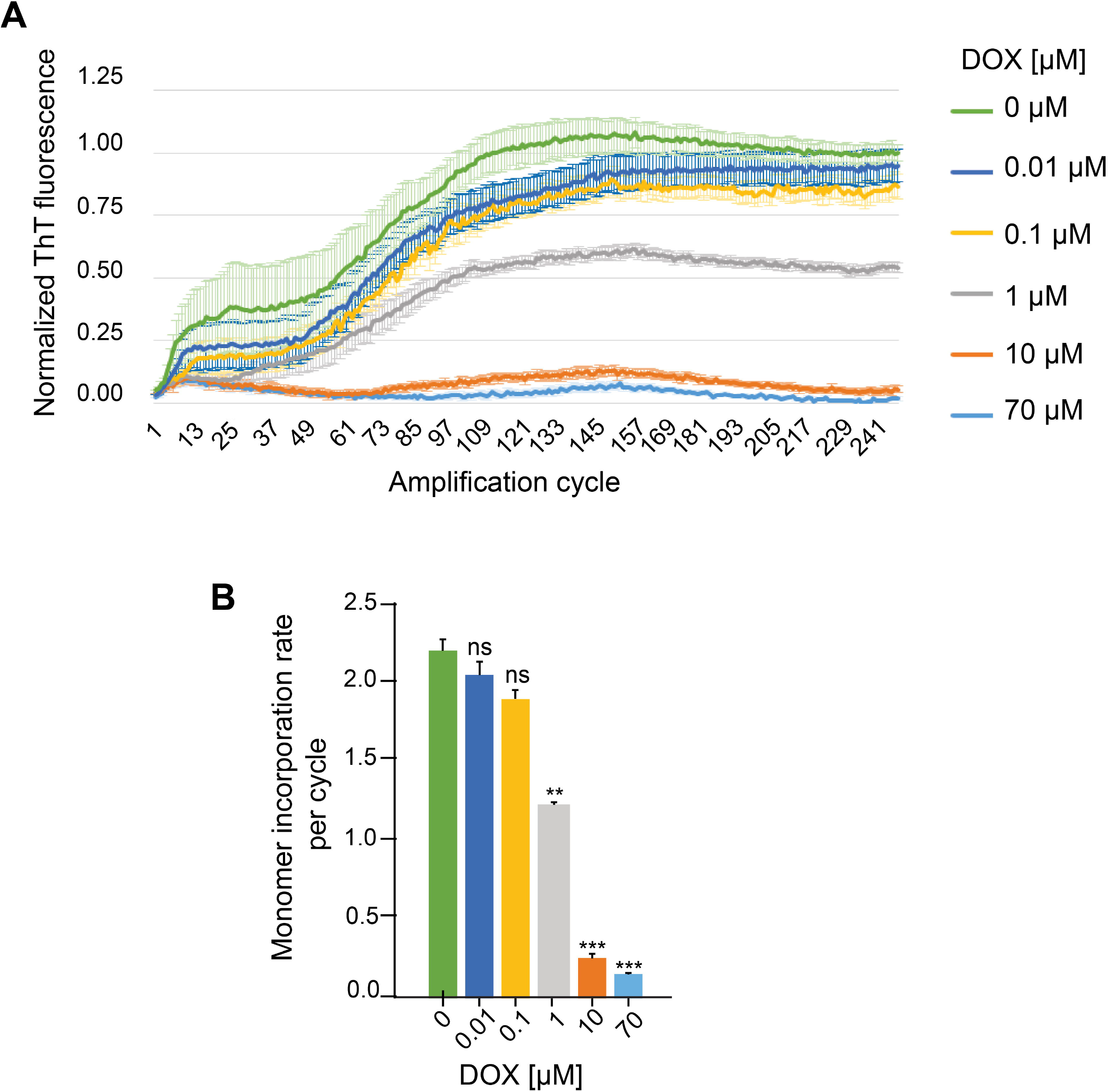
Effect of different doxycycline concentrations on *in vitro* amplification of K18 peptides by RT-QuiC. **A.** ThT fluorescence of 0.5 μM monomeric K18 incubated in the absence or presence of doxycycline. Three replicates of each sample were measured for 250 amplification cycles. **B.** Monomer incorporation rate per RT-QuiC cycle. n = 5 ns: not significant. * *p* < 0.05. ** *p* < 0.005. *** *p* <0.0005. Error bars represent SD.

## DISCUSSION

Recent results from our group and others [17,23,51–53] have shed light regarding the ability of tetracyclines in interfering with the toxicity and seeding of aggregate-prone proteins. In the present work, we extend these studies to tau. Our results demonstrate that doxycycline can disrupt tau aggregation in both heparin-induced and K18 self-assembling heparin-free systems (Fig. 1, 5 B and C). It is important to note that tau amyloid formation in the presence of doxycycline produces, in addition to oligomers, short fibrillary species (Fig. 4C). This effect is distinct from what has been previously reported for doxycycline on α-synuclein aggregation, where only oligomers are observed [23].

The surface hydrophobicity of tau aggregates has been associated with both cellular toxicity and their ability to serve as nucleation seeds for monomers [54]. Interaction of hydrophobic aggregated species with cell membranes, which induces permeability and ultimately dysfunction, is thought to be the mechanism responsible for toxicity [35,55,56]. Even inter-neuronal propagation of aggregated tau, often referred to as “spreading”, is governed by the hydrophobicity of pro-aggregant species [54]. Our results show that tau aggregates grown in the presence of doxycycline present less hydrophobic surfaces (Fig 2A), which could explain why tau species produced in the presence of this tetracycline are incapable of seeding (Fig. 2A) and ultimately trigger decreased toxicity in cell cultures (Fig. 2B).

Structural studies revealed that tau species produced in the presence of doxycycline have less β-structuration (Fig. 3C, F and G). This difference could ultimately affect the clearance of these species, since aggregates enriched in β-structures can directly or indirectly damage cellular proteostatic mechanisms, such as ubiquitin-proteasome (UPS), chaperone-mediated autophagy (CMA), and macroautophagy [57–61]. Therefore, the formation of tau species of smaller size (Fig. 4C) and lower /5-structure content (Fig. 3C, F and G) suggests that doxycycline could also indirectly benefit these proteostatic control systems.

The pattern of proteolytic digestion of species obtained in the absence or presence of doxycycline also revealed differences in the packing of the resultant fibrils (PHF-like and short filaments). Densitometric profiles revealed that digestion products of species formed in the presence of doxycycline were enriched in low molecular weight elements (Fig. 4 D and E). Trypsin limited digestion of PHFs *in vitro* results in a resistant fragment comprising the amyloid core, including approximately the second half of R1, R2, R3 and the first half of R4 [46]. Therefore, the absence of this 17 KDa trypsin resistant fragment in the mixture containing doxycycline suggests that the tetracycline might interact with the tau microtubule binding region which forms the amyloid core. The fact that doxycycline also inhibits K18 self-aggregation supports the hypothesis that the 4R pro-aggregant domain of tau is involved in doxycycline interaction. Moreover, these mechanisms observed *in vitro* could have impact in neuroprotection, since the presence of smaller aggregated species with higher susceptibility to proteases would also favor their “clearance” by the intracellular degradation systems. Likewise, taking into account that amyloid fibrils trigger microglia activation and chronic inflammatory response, which in turn lead to neurodegeneration [62–64], the doxycycline-increased susceptibility to proteases could be beneficial for neuron survival.

From a clinical perspective, doxycycline has been used for decades in human health and has proven to be a safe and well tolerated drug[65,66]. Moreover, due to its anti-inflammatory actions, doxycycline therapy is not restricted to the treatment of microbial infections, but has also been demonstrated to be useful for the management of periodontal and skin inflammatory pathologies [67,68]. When used as an antimicrobial compound, doxycycline (200 mg twice a day) achieves a concentration of about 3 μM in the cerebral spinal fluid (CSF) [69]. Therefore, considering its brain penetration, the tau/doxycycline ratio from our experiments, and the concentration of tau in CSF (approximately 244 pg/ml) [70], sub-antibiotic doses would be high enough to exert neuroprotection.

In addition, considering the concentration of doxycycline required for a protective effect, Umeda et al. [71] found, in a mouse model of AD, that older mice required higher concentrations of rifampicin to achieve a degree of protection comparable to younger mice. Therefore, antibiotic therapy could be especially valuable at the first stage of the disease. Likewise, considering the feasibility of doxycycline therapy for chronic treatments, as those required for these neurodegeneration, it is important to note that since sub-antibiotic doses would be high enough to reach neuroprotective concentrations in the brain, a selective pressure over native microbiota should not be imposed. Likewise, several reports found no evidence of microbiota disturbance during long-term doxycycline therapy even after two years of treatment with sub-antibiotic doses (20 mg/day) [72,73]. Interestingly, two sub-antibiotic doxycycline-formulations (Periostat and Oracea) have already been approved by de US FDA for long-term treatment of periodontal pathologies and the chronic inflammatory skin disease rosacea [72].

Regarding the benefits of doxycycline in neurology, daily doses of doxycycline and rifampicin during three month were found to significantly ameliorate dysfunctional behavior and cognitive decline in a clinical trial with more than one hundred patients with probable AD and mild dementia[74]. However, a later trial suggested that neither rifampicin nor doxycycline were capable of hindering the progression of neurodegeneration in diagnosed AD patients [75]. Since neurodegeneration involves the death of neurons leading to irreversible brain injury and, considering that amyloid aggregation is one of the first events of the deleterious cascade that takes place in tauopathies, it is plausible to think that drugs such as doxycycline that inhibit tau aggregation would be more suitable candidates for preventive rather than for palliative therapy. Therefore, the controversial results from clinical trials may be explained considering the administration time of the drug since once the appropriate staring points for treatment have passed its therapeutic action could decreased.

In summary, our results extend the spectrum of the anti-aggregation action of doxycycline over tau, a key player during the development and progression of tauopathies such as AD. Altogether, the anti-aggregant ability on tau reported herein, in addition to its long safety record, as well as its brain penetration and low concentration required, prompt us to propose doxycycline as a valuable candidate for the development of a therapy against Alzheimer’s disease and related tauopathies.

## Abbreviations

MAP: microtubule associated protein
IDP: intrinsically disordered protein
PHF: paired helical filaments
NFT: neurofibrillary tangle
AD: Alzheimer’s disease
DOX: doxycycline
PPII: type II polyproline helix.

## Acknowledgements

The authors would like to thank the Institut du Cerveau et de la Moelle Epinière (ICM), specially the electron microscopy team at ICM Quant Platform for assistance with microscopy services and the Cell Culture Platform. We also thank Mr. Claude Burgio and SkyBio for invaluable support. This work was supported by grants from PIP-CONICET 722, PICT-MINCYT3379, PICT2018-02989, PIUNT-UNT D644/1. LM, FGL and SS were supported by fellowships from the Argentinean Government (Becas Internas Doctorales y Posdoctoral from Consejo Nacional de Investigaciones Científicas y Técnicas). ADM is supported by a postdoctoral fellowship from the Galician Government (Programa de axuda á etapa posdoutoral, XUGA, GAIN, ED481B 2017/053). TFO is supported by the Deutsche Forschungsgemeinschaft (DFG, German Research Foundation) under Germany’s Excellence Strategy - EXC 2067/1-390729940.

